# Clinical and Cross-Domain Validation of an LLM-Guided, Literature-Based Gene Prioritization Framework

**DOI:** 10.64898/2026.01.22.701191

**Authors:** Taushif Khan, Ansarullah, Joshy George, Jeff Tomalka, Rafick P Sekaly, Karolina Palucka, Damien Chaussabel

**Affiliations:** The Jackson Laboratory of Genomic Medicine, Farmington, CT, 06032; The Jackson Laboratory, Bar Harbor, ME, 04605; Emory School of Medicine, Atlanta, GA 30307

## Abstract

**Background:** We previously published a literature-based pipeline for sepsis gene prioritization (PS3 and candidate genes) using an LLM-enabled retrieval and judging framework. Here, we extend that work to ask whether these prioritized genes show independent clinical validity and whether the same strategy generalizes to a drug–obesity– infection setting.

**Methods:** Using the original LLM-guided workflow, we evaluated PS3 and the Candidate set in two new settings. First, we tested 28-day mortality prediction in the independent VANISH sepsis trial, benchmarking PS3 and Candidate against two established immune signatures—the Severe-or-Mild (SoM) signature and Immune Health Metric (IHM)—under a uniform logistic-regression framework with clinical covariates. Second, we applied the same genome-wide screening and tiered judging pipeline to GLP-1/obesity/infection biology centered on semaglutide, comparing Tier-1 and Tier-2 gene sets to STEP trial serum proteomics at gene and Hallmark pathway levels. In parallel, we fine-tuned an open-weight GPT-OSS-20B model on curated sepsis justifications to obtain a domain-aware “LLM-as-judge,” and compared its scoring behavior with the base model on semaglutide Tier-2 genes.

**Results:** In the full VANISH cohort, PS3 and the Candidate set showed moderate discrimination, whereas SoM remained the strongest single predictor of 28-day mortality. In the Critical/High APACHE II subgroup, PS3 achieved ROC and precision–recall performance comparable to, or slightly better than, SoM despite its smaller, knowledge-derived composition, indicating that literature-prioritized genes capture mortality-relevant immune dysregulation under severe illness. In the semaglutide case study, gene-level overlap between LLM-prioritized genes and differentially abundant serum proteins was modest, but Tier-1 genes recapitulated the main semaglutide-responsive metabolic programs from STEP and highlighted additional immune–metabolic pathways relevant to infection, with discordances largely explained by serum proteome coverage. The fine-tuned judge remained moderately concordant with the base GPT-OSS across mechanistic themes, preserving overall ranking while inducing systematic, biologically interpretable shifts in immune and infection-related scores.

**Conclusions:** An LLM-guided, literature-based gene prioritization framework yields compact gene sets that show independent sepsis mortality signal and pathway-level concordance in a semaglutide/obesity/infection setting, while a sepsis-aware LLM-as-judge provides domain-specific refinements without overturning core rankings. Together, these findings support knowledge-grounded, LLM-derived gene sets and judges as interpretable components for probing immune dysregulation across diseases and therapies.

**Short abstract:** We previously published a literature-based pipeline for sepsis gene prioritization (Priority Set 3, PS3, and a Candidate set) using an LLM-enabled retrieval and judging framework; here, we test whether these prioritized genes show independent clinical validity and whether the same strategy generalizes to a drug–obesity–infection setting. Using the original workflow, we first evaluated 28-day mortality prediction in the independent VANISH sepsis trial, benchmarking PS3 and the Candidate set against two established immune signatures—the Severe-or-Mild (SoM) signature and Immune Health Metric (IHM)—within a uniform logistic-regression framework adjusted for clinical covariates. In the full cohort, PS3 and Candidate showed moderate discrimination, while SoM remained the strongest single predictor; however, in the Critical/High APACHE II subgroup, PS3 achieved ROC and precision–recall performance comparable to, or slightly better than, SoM despite its smaller, knowledge-driven composition, indicating that literature-prioritized genes capture mortality-relevant immune dysregulation under severe illness. We then applied the same genome-wide screening and tiered judging pipeline to GLP-1/obesity/infection biology centered on semaglutide, comparing Tier-1 and Tier-2 genes with STEP trial serum proteomics; although gene-level overlap with differentially abundant proteins was modest, Tier-1 genes recapitulated key semaglutide-responsive metabolic programs and highlighted additional immune–metabolic pathways relevant to infection, with discordances largely attributable to serum proteome coverage. Finally, supervised fine-tuning of an open-weight GPT-OSS model on curated sepsis justifications yielded a domain-aware “LLM-as-judge” that remained broadly concordant with the base model while inducing systematic, interpretable shifts in immune and infection-related scores. Together, these results support LLM-guided, literature-based gene sets and judges as compact, mechanistically interpretable components for probing immune dysregulation across diseases and therapies.

## Introduction

Sepsis remains a leading cause of morbidity and mortality in intensive care units worldwide and continues to challenge efforts toward precision risk stratification (Hotchkiss, Monneret and Payen 2013; Sweeney and Wong 2016; Chen *et al*. 2025). Although large-scale transcriptomic studies have revealed extensive immune dysregulation(Diorio *et al*. 2020; Zhang *et al*. 2023;), the translation of these findings into clinically actionable mortality predictors has been limited (Sweeney *et al*. 2018; Pelaia, Shojaei and McLean 2023; Moore *et al*. 2025). Many computational approaches rely on statistical feature selection from high-dimensional gene-expression data, producing signatures that often lack mechanistic grounding, vary across cohorts, and struggle to generalize in clinical settings (Wong *et al*. 2015; Davenport *et al*. 2016; Sweeney *et al*. 2018; Chenoweth *et al*. 2024). These challenges highlight a persistent gap between molecular signatures uncovered through data-driven discovery and the mechanistic understanding contained within decades of biomedical research.

To address this gap, we previously developed a knowledge-driven gene prioritization framework that uses large language models (LLMs) and retrieval-augmented evaluation to score genome-wide genes for their involvement in sepsis biology across multiple mechanistic dimensions (Khan *et al*. 2025). That work introduced Priority Set 3 (PS3) and a refined high-confidence Candidate set, both derived from structured, literature-grounded reasoning rather than direct optimization on clinical outcomes. While the original study demonstrated the biological coherence and evidence fidelity of the prioritization strategy, it did not assess the clinical utility of the resulting gene sets or how they behave inside a full retrieval-augmented generation (RAG) pipeline.

In this study, we extend that work in three directions. First, we systematically characterize the thematic landscape of the sepsis literature underlying our RAG. This embedding analysis allows us to visualize how gene- and pathway-centric queries land in the literature space and to show that retrieval choices, rather than the generative model alone, largely determine which mechanistic narratives are surfaced.

Second, we evaluate the ability of our previously published, literature-derived gene sets to predict 28-day mortality in an independent cohort from the VANISH trial(Antcliffe *et al*. 2019). This evaluation is intentionally comparative: we benchmark PS3 and the Candidate set against two well-established, data-driven immune signatures—the Severe-or-Mild (SoM) signature(Zheng *et al*. 2021; Ganesan *et al*. 2025) and the Immune Health Metric (IHM)(Sparks *et al*. 2024). These signatures have been validated across diverse infectious and inflammatory conditions and represent high-performing baselines for assessing mortality-associated immune states. By examining performance across the full cohort and severity-stratified subgroups, we ask whether literature-grounded prioritization recovers mortality-relevant immune dysregulation and whether any signal strengthens under critical illness, where risk is highest.

Third, we test the cross-disease portability of this framework using semaglutide, a GLP-1 receptor agonist with proven efficacy in obesity and type 2 diabetes (Yugar *et al*. 2024) and emerging evidence for broader cardiometabolic and inflammatory effects (Taktaz *et al*. 2024). We apply the same LLM-based screening pipeline to identify GLP-1/obesity/infection genes and compare these to serum proteomic changes observed in a large randomized trial of semaglutide in obesity (Maretty *et al*. 2025). Although gene-level overlap with differentially abundant proteins is modest, we show that pathway-level concordance is stronger: the prioritized genes are enriched for the same metabolic, inflammatory, and vascular programs perturbed by semaglutide and highlight a subset of candidates with plausible roles in latent or chronic infection biology.

A final aim is to explore whether high-confidence, citation-supported outputs from our prior pipeline can be repurposed to align an open-weight LLM for scientific evaluation. General-purpose LLMs can summarize biomedical text but often exhibit inconsistent citation behavior, variable reasoning structure (Kontsioti, Maskell and Pirmohamed 2024; Qiu *et al*. 2025), and occasional hallucinations (Farquhar *et al*. 2024; Asgari *et al*. 2025; Omar *et al*. 2025; Shool *et al*. 2025). To address these limitations, we curated mechanistic justifications from the sepsis prioritization experiments and used them to supervise fine-tuning of a 20B-parameter model (GPT-OSS-20B) (OpenAI *et al*. 2025). The resulting model is intended to function as an “LLM-as-a-Judge”: a domain-focused evaluator that reads retrieved evidence, generates structured mechanistic reasoning, and cites supporting PMIDs transparently.

Together, these components—literature-space mapping, mortality prediction, cross-domain semaglutide application, and LLM-as-a-Judge development—provide an integrated evaluation of both the predictive and interpretive value of literature-driven gene prioritization. Our results show that knowledge-grounded gene sets capture mortality-associated immune disruption comparable to established signatures in the sickest patients, that the same framework generalizes to GLP-1 biology with stronger agreement at the pathway than at the single-gene level, and that curated justifications can align an open-weight model toward more structured, citation-aware scientific judgment while leaving retrieval quality as a dominant constraint. This work outlines a generalizable template for building and stress-testing knowledge-grounded biomarker frameworks across diseases.

## Methods

### Literature embedding and cluster-level context annotation

To convert unstructured sepsis literature into structured retrieval contexts for our RAG system, we constructed a document-level embedding space from PubMed records related to sepsis(Khan *et al*. 2025). Records were segmented into 191,570 text nodes (abstracts or shorter chunks), each embedded with SPECTER2 (Singh *et al*. 2022) to obtain 768-dimensional vectors. We applied principal component analysis (PCA) for initial dimensionality reduction and Uniform Manifold Approximation and Projection (UMAP) for 2-dimensional visualization. A k-nearest-neighbor graph built on the high-dimensional embeddings was partitioned using the Leiden community detection algorithm, yielding 15 meta-clusters that define broad thematic “contexts” within the knowledge base.

For each cluster, we used Claude Sonnet-4 to generate structured annotations summarizing (i) the primary topic, (ii) clinical relevance, and (iii) predominant study designs. Annotations were stored in a JSON schema and used to interpret which regions of the embedding space were accessed by RAG queries.

### Gene Sets Evaluated

We evaluated gene signatures generated in our previously published literature-driven prioritization framework, including Priority Set 3 (PS3) and a refined high-confidence Candidate set (Table S1). These signatures were derived through structured assessment of genome-wide transcripts across multiple dimensions of sepsis biology, integrating mechanistic, immunologic, biomarker, and therapeutic evidence from the biomedical literature. For benchmarking, we included two established immune response signatures: the Severe-or-Mild (SoM) signature (Zheng *et al*. 2021; Ganesan *et al*. 2025), representing a conserved dysregulated immune state linked to poor outcomes across infectious diseases, and the Immune Health Metric (IHM), a transcriptomic signature quantifying deviation from immune homeostasis (Sparks *et al*. 2024). Complete gene lists for all four signatures are provided in the Supplementary Materials (Table S1).

### VANISH Transcriptomic Preprocessing and Mortality Prediction Modeling

Whole-blood gene-expression profiles from the VANISH randomized trial were generated by genome-wide microarray profiling of RNA extracted from peripheral blood collected at enrollment, as previously described (Antcliffe *et al*. 2019; Ganesan *et al*. 2025). Raw microarray intensities were log_2_-transformed and normalized using the established preprocessing workflow for this cohort. For each gene-set analysis, we restricted predictors to genes belonging to the corresponding signature. Continuous variables—including gene expression values, age, and APACHE II score—were standardized to zero mean and unit variance within each training fold. Missing data were rare and were imputed within folds using the median for continuous variables and the mode for categorical variables to avoid information leakage.

To assess the prognostic utility of each gene set, we fit multivariable logistic regression models that combined the selected gene-expression features with the three clinical covariates (age, sex, APACHE II score). The same modeling strategy was applied to all signatures to allow direct comparison. Predictive performance was estimated using stratified five-fold cross-validation, preserving the proportion of deaths in each fold. For every fold we computed the area under the receiver operating characteristic curve (ROC-AUC), area under the precision–recall curve (PR-AUC), overall accuracy, and the positive predictive value at 90% sensitivity. Metrics were averaged across folds to obtain stable estimates. Differences in ROC-AUC between gene sets were evaluated using DeLong’s test, and accuracy differences were assessed via permutation resampling of fold assignments. Stability of logistic-regression coefficients across folds was examined to support interpretability of model estimates.

### WideNet Screening and Tiered Gene-Set Selection for GLP-1/Semaglutide

We applied our genome-wide LLM screening framework (WideNet) to all human protein-coding genes, issuing structured prompts that asked, for each gene, whether there was evidence for involvement in (i) obesity and metabolic regulation (body weight, adiposity, insulin resistance, lipid metabolism), (ii) infection and host defence (innate/adaptive immunity, chronic or latent infection), and (iii) GLP-1–related mechanisms. For each axis, GPT-4o returned a numerical association score from 0–10 together with a short free-text rationale. Using the same scoring rubric as in our sepsis analyses (Toufiq *et al*. 2023; Khan *et al*. 2025), we retained genes with a GLP-1–axis score ≥6, yielding a Tier-1 set of 1,419 GLP-1/obesity/infection–associated genes aggregated from the >19 genes queried per question in the Tier-1 prompt set (scores of Tier1 and Tier 2 Table S2.1).

Within Tier 1, we then performed a second, semaglutide-focused pass. For each gene we asked whether semaglutide (or semaglutide-anchored GLP-1RA trials) modulates that gene, its encoded protein, or its proximal pathway in humans with obesity or type 2 diabetes, and whether this link was supported by trial data, proteomics, or mechanistic studies versus being primarily inferential. These semaglutide-specific queries were scored from 0–10 using GPT-5. Genes that were both highly scored on the GLP-1 axis and received strong semaglutide-specific scores formed the Tier-2 set of 69 “directly semaglutide-linked” genes. This Tier-2 set was then subjected to retrieval-augmented generation (RAG) and hybrid LLM evaluation using the same semaglutide-association prompt set as in Tier 2. Full prompt templates, scoring rubrics, and decision rules for Tier-1 and Tier-2 gene selection are provided in the Supplementary Methods.

### Proteomic comparison set and evaluation

As an external reference for semaglutide-responsive biology, we used serum proteomics from the STEP semaglutide versus placebo trials, which quantified ∼6,400 circulating proteins using SomaScan technology (Maretty *et al*. 2025). We mapped Tier-1 and Tier-2 genes to this panel via their encoded proteins and defined “semaglutide-responsive” proteins as those reported as significantly different between semaglutide and placebo in the original analysis after Holm–Bonferroni correction. At the gene level, we counted overlaps between our LLM-derived Tier-1 and Tier-2 gene sets and these semaglutide-responsive proteins.

For pathway-level validation, we performed enrichment analysis with a GSEA-style over-representation framework (enricher) using the HALLMARK gene sets from MSigDB (Liberzon *et al*. 2015; Kuleshov *et al*. 2016; Fang, Liu and Peltz 2023) . Tier-1 GLP-1/obesity/infection genes were tested for enrichment against a background of all proteins in the STEP panel and, in a focused analysis, against the subset of semaglutide-responsive proteins. Enriched HALLMARK pathways from the Tier-1 gene set were then compared with the HALLMARK programmes reported as altered by semaglutide in the STEP proteomics (for example, xenobiotic and fatty-acid metabolism, mTORC1 signalling, glycolysis, cholesterol homeostasis, adipogenesis, bile-acid metabolism and immune/inflammatory pathways).

### Supervised fine-tuning of a sepsis-aware GPT-OSS-20B LLM-as-judge

We fine-tuned the open-weight GPT-OSS-20B causal language model to align its reasoning with sepsis-specific literature patterns. Because long-context training (∼3,072 tokens) and stable optimization were required, we performed full-parameter supervised fine-tuning rather than adapter-based approaches. Training followed an answer-only causal language modeling objective, with all instruction and input tokens masked in the loss function. Optimization used AdamW with a learning rate of 1×10^−5^, cosine learning-rate scheduling, an effective batch size of 16 achieved through gradient accumulation, and bfloat16 precision. Gradient checkpointing was applied for memory efficiency. Models were trained for three epochs on a multi-GPU high-performance computing node equipped with NVIDIA V100/A100 GPUs. Checkpoints were evaluated after each epoch, and the final checkpoint was selected based on minimum validation loss.

Model performance was first assessed using intrinsic metrics (validation and test loss, perplexity) and qualitative review of a randomly sampled subset of outputs for factual accuracy, linkage between mechanistic claims and citations, and adherence to the sepsis-specific task structure. To test whether the fine-tuned model could act as a generalizable “LLM judge,” we then evaluated it on an out-of-context task using semaglutide-related prompts and responses, comparing its scoring and prioritization behavior directly with the untuned GPT-OSS-20B under identical prompting conditions.

### Large language models used and deployment settings

We used a mix of frontier API models and locally hosted open-weight models for different stages of the pipeline (literature annotation, genome-wide screening, and judging). Frontier models were accessed via the OpenAI API and included **gpt-4o, gpt-5**, and **gpt-5-mini**, which were used primarily for high-precision tasks such as genome-wide gene screening prompts (“WideNet”), semaglutide/GLP-1–focused queries, and structured justification generation.

Open-weight models were deployed locally via Ollama and Hugging Face backends and included Llama 3.2, Phi-4, and GPT-OSS-20B. Among these, GPT-OSS-20B served as the base model for our supervised fine-tuned LLM-as-judge, while Llama 3.2 and Phi-4 were used for exploratory screening and ablation experiments. All models were run in strictly retrieval-augmented settings for sepsis-related tasks, with explicit instructions to ground answers in the provided literature snippets and to surface PubMed identifiers when available.

Claude Sonnet-4 was used specifically for meta-cluster annotation in the embedding analysis, where it generated structured summaries of cluster themes, clinical relevance, and study designs based on representative node subsets.

## Results

### From Unstructured Text to Structured Knowledge Contexts

The 15 Leiden meta-clusters derived from the literature embedding span a broad and interpretable spectrum of sepsis research (Table S2, Figure 1A). One group of clusters (1–4) capture core acute pathophysiology, including autonomic and cardiovascular regulation, neutrophil-driven inflammation, endothelial barrier dysfunction, and sepsis-associated acute kidney injury, each linked to characteristic experimental or clinical study designs. A second set of clusters (7, 8, 9, 10, 12–14) map distinct clinical contexts and vulnerable populations-for example HIV-associated and other immunocompromised states, T-cell– mediated immune dysfunction, long-term neuromuscular sequelae, pediatric and neonatal sepsis, nutritional interventions, and transplantation with pharmacologic immunosuppression. Additional clusters emphasize molecular mechanisms and patient heterogeneity: cluster 13 focuses on molecular biology and gene regulation relevant to biomarker and therapeutic target discovery, whereas cluster 15 concentrates on host genetics and personalized medicine. In contrast, clusters 5, 6 and 11 represent structural components of articles—reference lists, tables and figure captions, and discussion sections—as indicated by section headers and citation patterns. Together, these patterns show that the embedding-based vector database separates formal document structure from biological and clinical content, effectively converting unstructured sepsis texts into a set of well-defined, clinically meaningful “contexts” that can be selectively addressed by our RAG system.

**Figure 1.**
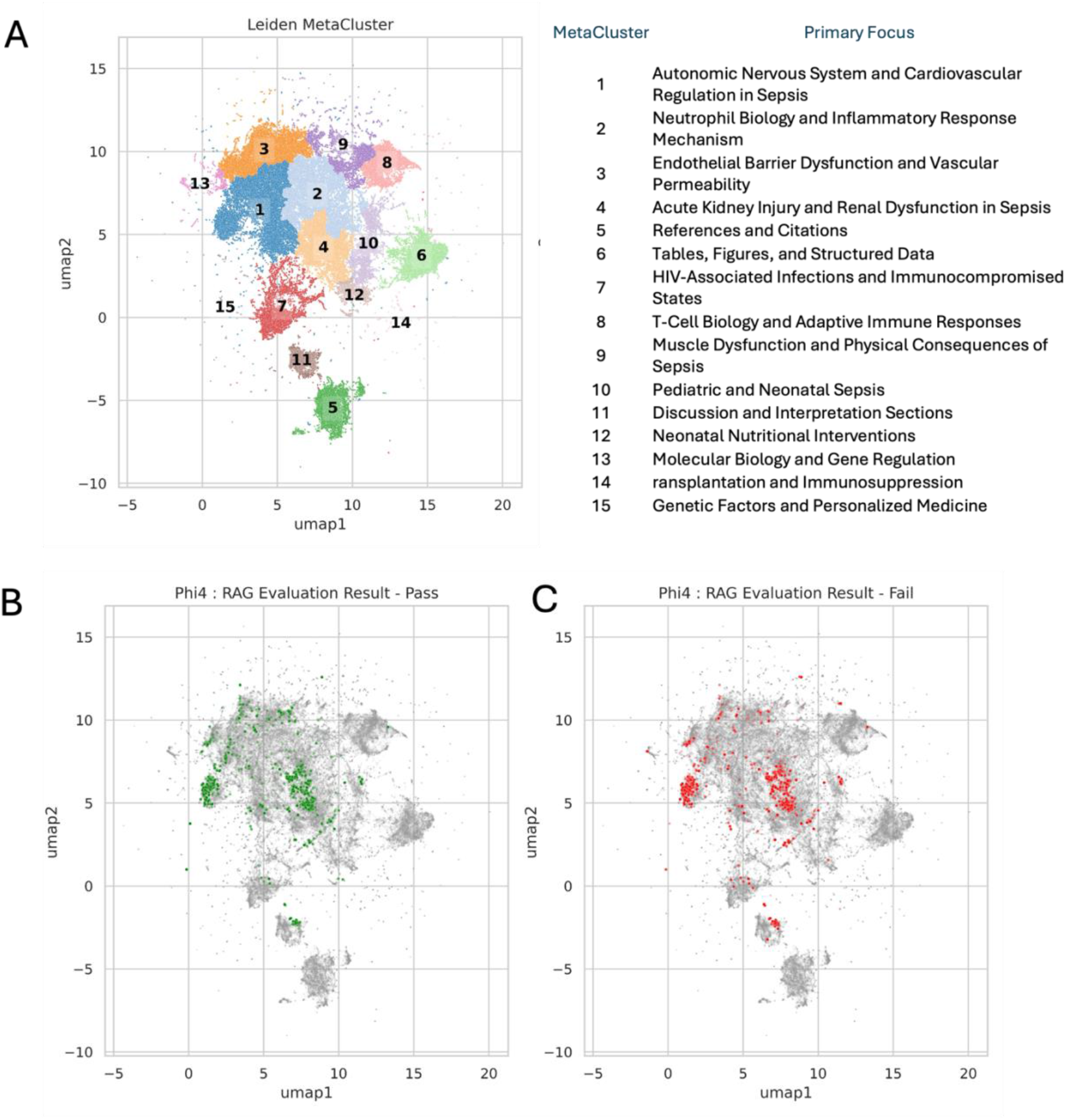
Structure of the sepsis literature embedding space and coverage by RAG retrieval and faithfulness assessment. SPECTER2 embeddings of 191,570 sepsis-related text nodes were projected to two dimensions using PCA followed by UMAP and clustered with the Leiden algorithm. **(A)** UMAP representation of the full corpus, with each point representing a document node and colors indicating the 15 Leiden meta-clusters (numbered 1–15); cluster-level primary focus, clinical relevance, and predominant methodology are summarized in Table S2. **(B**) in green, nodes that were selected as part of the top-5 reranked retrieval set for at least one of 4,872 RAG queries and whose evidence was judged *faithful* (pass) by the general-purpose LLM evaluator; all other nodes in the corpus are shown in grey.**(C)** In red, nodes that were retrieved in the top-5 but whose evidence *failed* the faithfulness check, again plotted over the full corpus in grey. The similar spatial distribution of pass and fail nodes indicates that current queries repeatedly sample a limited set of regions in the embedding space, while large areas of the sepsis knowledge graph remain unexplored by the present RAG configuration.

The resulting embedding space contained 191,570 unique literature nodes. Across 4,872 evaluation queries, the RAG pipeline drew evidence from 52,704 node–query pairings, corresponding to only 2,877 unique nodes (∼1.5% of the available corpus). Thus, while individual nodes were reused frequently as context, most of the literature space remained untouched by the current query set. When overlaid on the UMAP representation, nodes whose evidence passed the faithfulness check and those that failed exhibited broadly similar spatial distributions, despite all being selected from reranked top-5 retrievals. This pattern is consistent with the idea that the retrieval stage primarily determines where in the thematic landscape queries land, and that both faithful and unfaithful answers can arise from similar regions of the literature (Figure 1B, C). Overall, this analysis demonstrates that large-scale sepsis literature can be organized into interpretable, reusable knowledge contexts for LLM-based reasoning. In the next section, we move from mapping the literature space to testing a concrete example of knowledge-driven curation: a previously published priority gene set (PS3) evaluated for mortality prediction in an independent sepsis cohort.

### Evaluation of Knowledge-Derived Gene Sets for Sepsis Mortality Prediction

Predictive modeling using logistic regression revealed distinct performance patterns across the four gene sets (Table S1, Figure 2, S2). In the full VANISH cohort (Task 1; n = 175), the SoM gene set achieved the highest ROC-AUC for 28-day mortality (0.78 [0.61–0.91]) and strong precision–recall performance (PR-AUC = 0.82), along with the highest positive predictive value (PPV = 0.76) at 90% sensitivity (Table S3). The knowledge-derived PS3 and Candidate gene sets showed more modest ROC-AUC values (0.67 and 0.62, respectively) but maintained competitive precision (PR-AUC = 0.87 and 0.67) (Figure S2). Notably, the PR-AUC for PS3 slightly exceeded that of SoM despite its lower ROC-AUC, indicating that this more compact, literature-based signature retains good precision at comparable levels of recall. Although the IHM signature exhibited the highest overall accuracy (0.92), its lower ROC-AUC (0.61) suggested limited discriminative ability in this heterogeneous cohort, consistent with the expectation that high accuracy can arise in imbalanced datasets even when class separation is suboptimal.

**Figure 2.**
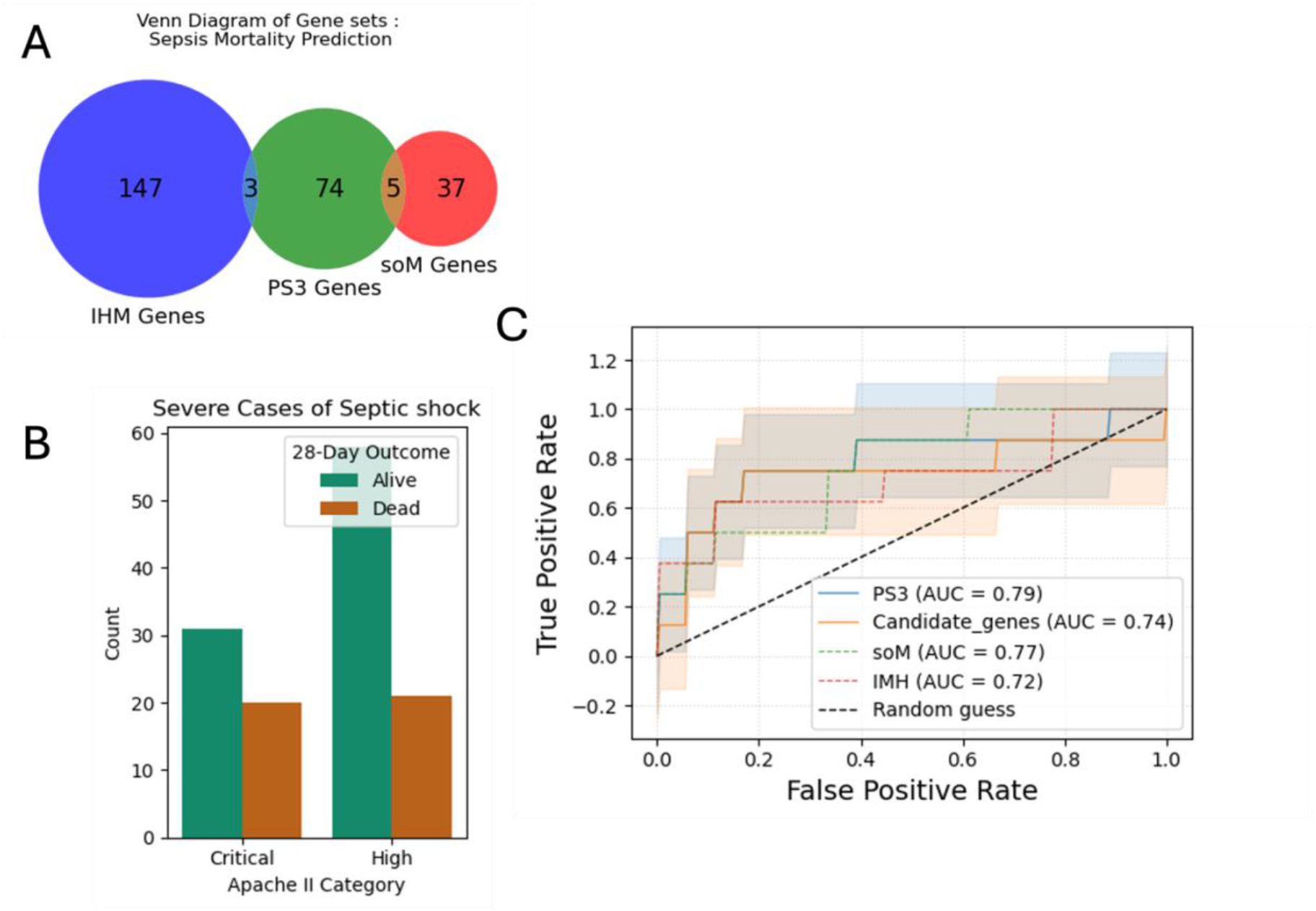
Overlap of gene sets and mortality prediction performance in severe sepsis. **(A)** Venn diagram comparing the three gene sets evaluated for 28-day mortality prediction: the Immune Health Metric (IHM) signature, the LLM-prioritized Priority Set 3 (PS3) genes, and the Severe-or-Mild (SoM) signature. Most genes in each set are unique (147 IHM, 74 PS3, 37 SoM), with only a small overlap between PS3 and IHM (3 genes) and between PS3 and SoM (5 genes), indicating that PS3 captures largely distinct biology from existing immune signatures. **(B)** Distribution of 28-day outcomes among VANISH patients with severe septic shock (Critical and High APACHE II categories) used for the restricted mortality analysis, showing the number of survivors versus non-survivors in each severity stratum and the resulting class imbalance. **(C)** Receiver-operating characteristic curves for 28-day mortality in this severe subgroup from logistic-regression models based on each gene set (with clinical covariates). PS3 achieves the highest discrimination (AUC = 0.79), followed by SoM (AUC = 0.77), the Candidate gene set (AUC = 0.74) and IHM (AUC = 0.72), all outperforming random guessing, showing that a literature-driven, LLM-prioritized signature can match or modestly exceed established data-driven immune signatures in the highest-risk patients.

When analysis was restricted to patients with Critical or High APACHE II scores (Task 2; n = 130), performance across signatures converged and the relative ranking shifted (Figure 2, Table S3). In this high-risk subgroup, PS3 achieved the highest ROC-AUC (0.79 [0.53–0.97]) and PR-AUC (0.90), marginally outperforming SoM (ROC-AUC = 0.77 [0.55–0.93]; PR-AUC ≈ 0.90) and maintaining a significant advantage over SoM under FDR-adjusted comparison (PS3 vs SoM, p ≈ 0.003). While SoM preserved higher precision at high sensitivity (PPV = 0.78 vs 0.65 for PS3 at matched specificity and sensitivity thresholds), the near-identical PR-AUCs indicate that both signatures sustain strong precision–recall performance under severe disease conditions. The Candidate set showed intermediate results (ROC-AUC = 0.74 [0.47–0.97]; PR-AUC = 0.71), whereas IHM again displayed high overall accuracy (0.95) but lower discrimination (ROC-AUC = 0.72). Pairwise ROC comparisons using DeLong’s test (Supplementary Table S4) showed no significant difference between PS3 and SoM in either the full cohort (p = 0.64) or the Critical/High APACHE subgroup (p = 0.88). By contrast, Candidate and IHM differed significantly from the other signatures in both settings (all p < 0.01), consistent with their lower ROC-AUC estimates. Together, these findings indicate that the knowledge-grounded PS3 set, although smaller and literature-derived, performs comparably to the data-driven SoM signature for mortality discrimination in high-risk patients.

### Semaglutide / obesity / infection case study

To test whether our LLM-based genome-wide screening and judging framework generalizes beyond sepsis, we applied it to semaglutide, a GLP-1 receptor agonist with proven efficacy for obesity and type 2 diabetes and emerging evidence for broad cardiometabolic and inflammatory benefits. We specifically asked whether the same pipeline could (i) retrieve genes and pathways implicated in semaglutide response in obesity and (ii) highlight genes with plausible roles in modulating latent or chronic infection.

Across the 6,386 proteins quantified in the STEP serum proteomics, our Tier-1 GLP-1/obesity/infection screen (≈1,400 genes) mapped 790 genes to measured proteins, of which 83 showed significant semaglutide–placebo differences (Table S3); in Tier-2, 53 of the 69 semaglutide-focused genes were measurable and 8 of these passed the same proteomic filter (Table S4, Figure S3C). While gene-level overlaps with STEP proteomics were modest (Figure S3C), the pathway-level view revealed a more nuanced pattern (Figure 3A). Using a common background of STEP-measurable proteins, the Nature Medicine semaglutide-responsive gene set was significantly enriched for Hallmark xenobiotic metabolism, fatty-acid metabolism, coagulation, epithelial–mesenchymal transition and complement pathways (Figure 3, Table S5). The Tier-1 GLP-1/obesity/infection screen partially recapitulated this footprint, converging on xenobiotic metabolism, coagulation, epithelial– mesenchymal transition and complement but not on fatty-acid metabolism. By contrast, the Tier-2 semaglutide-focused genes showed limited overlap with the Nature Medicine profile, aligning only on complement, but were strongly enriched for pathways that did not reach significance in the original STEP analysis, including pancreatic β cell programs and multiple inflammatory/immune axes (e.g. TNFα/NF-κB, IL-6/JAK/STAT3, interferon responses and other stress/injury signatures). Thus, the LLM-derived Tier-1 and Tier-2 gene sets recover part of the known semaglutide metabolic signature while emphasizing additional β cell and inflammatory components that may be more tightly linked to GLP-1 biology and infection risk.

**Figure 3.**
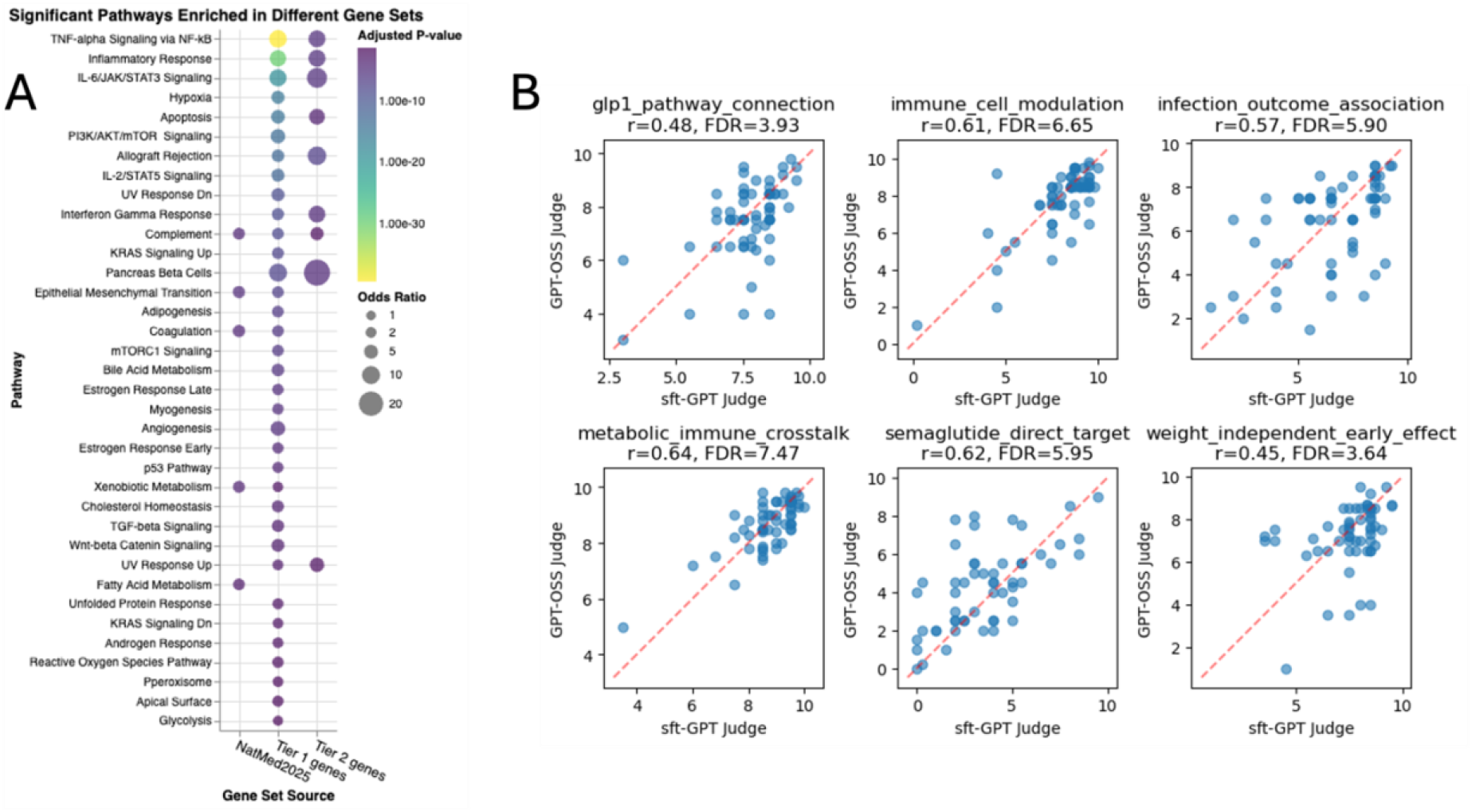
**A. Hallmark pathway enrichment across semaglutide-responsive and LLM-derived gene sets.** Dot plot showing Hallmark (MSigDB) pathways that were significantly enriched for at least one of three gene sets: semaglutide-responsive proteins from the STEP Nature Medicine study (“NatMed2025 genes”), the Tier-1 GLP-1/obesity/infection screen, and the Tier-2 semaglutide-focused genes. Enrichment was assessed by over-representation analysis using a common background of proteins measurable in the STEP proteomics panel, with Benjamini–Hochberg–adjusted p-values. Dot size encodes the odds ratio for each pathway– gene set combination, and dot color represents the adjusted p-value (darker points indicate stronger enrichment). **B. Concordance between base and fine-tuned LLM judges on semaglutide-related genes**. Scatter plots compare theme-specific scores assigned by the base GPT-OSS judge (y-axis) and the sepsis-tuned SFT-GPT-OSS judge (x-axis) for 69 high-priority semaglutide/GLP1-related genes. Each point represents a single gene. Panels show six predefined biological themes: GLP1 pathway connection, immune cell modulation, infection outcome association, metabolic–immune crosstalk, semaglutide direct target, and weight-independent early effect. The red dashed line denotes the line of identity (y = x), where both judges assign identical scores. Because scores are used primarily in a rank-like manner, agreement between judges is summarized using Spearman rank correlation (ρ) with false-discovery-rate–adjusted p-values (FDR) reported in each panel.

Notably, canonical GLP-1 pathway genes such as GLP1R, GCG and PRKACA were consistently flagged by the LLM–RAG–Judge as semaglutide-associated but showed no significant change in circulating protein levels (Maretty *et al*. 2025). This pattern is expected for membrane and intracellular signaling proteins, which are poorly sampled by serum proteomics. It reflects a mismatch between a gene-wise mechanistic task and a serum-only proteomic readout, rather than a failure of the literature-based prioritization. Taken together, these results suggest that agreement between our hybrid LLM platform and STEP proteomics is most naturally interpreted at the pathway level, with gene-level discordances largely attributable to assay coverage and compartment effects rather than solely to model error.

### Fine-tuning a sepsis-aware LLM-as-judge and application to Tier 2 semaglutide genes

To obtain a domain-aware LLM-as-judge, we performed supervised fine-tuning (SFT) of GPT-OSS on the curated “sepsis” question–answer pairs used in our earlier experiments. Training was run for three epochs with a maximum context of 3,072 tokens and learning rate warm-up followed by linear decay (Table S7). Training logs showed a progressive reduction in loss from epoch 2.14 to 2.50 (0.147 → 0.130) with stable gradient norms (∼7–7.5) as the learning rate decreased from 2.1×10^−6^ to 7.7×10^−7^. Between epochs 2.50 and 2.86 the loss plateaued around 0.13–0.14, after which the final logged step at epoch 3 showed an abrupt increase in loss (0.39). We therefore selected the best checkpoint obtained just before this deterioration (≈2.5–2.8 epochs) hereafter referred to as SFT-GPT-OSS. This pattern suggests that the model had already reached a favorable operating region prior to the end of the third epoch and that the final step reflects training instability rather than further improvement.

The fine-tuned model plays a purely evaluative role: it does not generate candidate genes but instead scores and ranks gene–mechanism statements produced by a separate “task LLM.” In our pipeline, Llama-3.2 generates structured justifications for each gene, and either the base GPT-OSS or SFT-GPT-OSS acts as the judge that assigns theme-specific scores. This design isolates the effect of fine-tuning on the *judging* step while holding the upstream proposal step constant.

#### Concordance of semaglutide gene scores between base and fine-tuned judges

We next asked how SFT changes the behavior of the judge in a realistic, non-sepsis use case. As a test bed we used the 69 Tier 2 genes that our hybrid LLM platform had previously linked to semaglutide or GLP1R biology (Figure 3B). For each gene, Llama-3.2 produced explanations and numerical scores for six predefined biological themes: “GLP1 pathway connection,” “immune cell modulation,” “infection outcome association,” “metabolic– immune crosstalk,” “semaglutide direct target,” and “weight-independent early effects.” These same prompts and rationales were then independently scored by the base GPT-OSS judge and the fine-tuned SFT-GPT-OSS judge, and their scores were compared using Spearman rank correlation (ρ) (Figure 3B), reflecting the rank-like use of scores in our downstream pipeline.

Across all six themes, SFT-GPT-OSS remained moderately concordant with the original GPT-OSS judge (ρ ≈ 0.45–0.64). Agreement was strongest for “metabolic–immune crosstalk” and “semaglutide direct target” (ρ ≈ 0.62–0.64), followed by “immune cell modulation” and “infection outcome association” (ρ ≈ 0.57–0.61). Even for “GLP1 pathway connection” and “weight-independent early effects,” which showed slightly broader dispersion, the majority of genes clustered near the identity line. False-discovery-rate–adjusted p-values were below 0.001 for all six themes, indicating that the observed concordance is unlikely to arise by chance in this exploratory setting.

Taken together, these analyses show that SFT on sepsis-specific judgments does not dramatically re-order semaglutide-related genes, but instead induces modest, systematic shifts while preserving the overall ranking structure. In practice, the fine-tuned judge behaves as a sepsis-informed variant of GPT-OSS that maintains global scoring behavior yet can subtly reweight evidence in mechanistic themes—particularly those involving immune and infection-related biology—without destabilizing the underlying prioritization. However, because our evaluation is cross-domain and lacks an external gold-standard ranking, these results are best interpreted as behavioral characterization rather than definitive proof that SFT-GPT-OSS is superior to the base judge for semaglutide-focused tasks.

## Discussion

Using a uniform modeling strategy across four gene sets, we found that an LLM-guided, knowledge-driven signature can approach—though not consistently surpass—the performance of established data-driven immune signatures for sepsis mortality prediction (Figure 2, Table S3). In the overall VANISH cohort, the SoM signature remained the strongest single predictor, with the highest AUC and best PPV at 90 % sensitivity, consistent with its design as a broadly applicable marker of systemic immune dysregulation across infections and risk factors. By contrast, PS3 and the Candidate set—derived from literature-based evaluation of sepsis mechanisms rather than direct optimization on outcome data—showed only modest discrimination in this mixed-severity setting. However, when analysis was restricted to patients with Critical and High APACHE II scores, PS3 achieved the highest ROC AUC and PR AUC and tracked closely with SoM, suggesting that the LLM-prioritized genes do capture mortality-relevant signal that becomes most apparent under conditions of pronounced immune and organ dysfunction.

These findings fit with the complexity and breadth of the host immune signatures captured by SoM (Zheng *et al*. 2021; Ganesan *et al*. 2025) and IHM(Sparks *et al*. 2024). SoM was trained on more than 12,000 blood transcriptomes and proteomes from diverse infections and clinical contexts, and IHM defines a quantitative “immune health” axis spanning a wide range of immune perturbations. Both therefore function as broad, high-capacity summaries of immune deviation, and it is reassuring that in our analysis they show stable performance across tasks. The observation that PS3—constructed purely from a knowledge-driven LLM+RAG pipeline—can match or slightly outperform these signatures in the sickest patients supports the idea that literature-prioritized genes can recover core elements of sepsis-specific immune derangement, even without being tuned on VANISH outcomes. At the same time, the relative underperformance of PS3 and the Candidate set in the full cohort highlights that our current prompting strategy, which emphasized general sepsis biology (pathogenesis, pathways, organ dysfunction, biomarkers, therapeutics) rather than outcome-specific evidence, is not yet optimized for mortality prediction.

The semaglutide case study provides a modest but informative test of whether our LLM-based screening and judging framework can be applied outside sepsis. Starting from GLP-1/obesity/infection prompts, the Tier-1 gene set showed limited gene-level overlap with semaglutide-responsive proteins in STEP but did recapitulate the main metabolic Hallmark program reported in the original proteomic analysis and additionally highlighted immunometabolic pathways linked to inflammation and host defense (Figure 3A). We view this primarily as pathway-level concordance, not strong validation of specific genes, especially because canonical GLP-1 signaling components (e.g. GLP1R, GCG, PRKACA) were prioritized by the LLM yet showed no detectable change in circulating protein levels, which is consistent with a fasting serum assay that incompletely captures tissue-localized signaling. More broadly, the case study underscores that a gene-wise, mechanism-focused LLM task and serum proteomics offer complementary but non-identical snapshots of drug biology, and that conclusions here should be treated as illustrative and hypothesis-generating rather than definitive.

Finally, fine-tuning GPT-OSS on curated sepsis question–answer pairs produced a sepsis-aware LLM-as-judge that behaves as a calibrated variant of the base model rather than a qualitatively new evaluator. In the semaglutide Tier-2 analysis, the fine-tuned judge remained moderately concordant with the original GPT-OSS across all mechanistic themes (Spearman ρ ≈ 0.45–0.64), indicating that domain-specific SFT largely preserves the global ranking of gene–mechanism statements while allowing modest, systematic shifts—most evident for immune and infection-related dimensions (Figure 3B). This pattern aligns with emerging evidence that LLM judges can be specialized for domains without losing general evaluative capacity, and that such judges can approximate expert assessments at scale. However, our analysis is cross-domain, confined to a single fine-tuned model, and lacks a gold-standard semaglutide ranking. We therefore interpret these findings as a cautious demonstration that sepsis-specific SFT can reshape an LLM judge’s behavior in biologically interpretable ways, rather than as evidence that SFT-GPT-OSS is uniformly superior to the base judge for GLP-1–related or other applications.

Taken together, these results argue less for building a single mortality-tuned signature and more for using LLM-guided prioritization as a flexible way to explore immune dysregulation across contexts. Rather than immediately retuning the pipeline around survival endpoints, a natural next step is to broaden the prompting and evaluation framework to capture shared axes of immune activation, resolution, and tissue injury that recur in sepsis, chronic cardiometabolic disease, and obesity-induced inflammation. Within such a framework, SoM and IHM can serve as anchoring “immune state” references, while LLM-derived gene sets define more focused mechanistic modules that can be followed across domains—for example, tracking how neutrophil, endothelial, and metabolic-immune programmes shift from acute sepsis to obesity and semaglutide response. The fine-tuned LLM-as-judge then becomes a reusable component for scoring these modules and for probing how domain-specific SFT (e.g., sepsis-, obesity-, or infection-focused) reshapes mechanistic interpretations. In this exploratory phase, the emphasis is on mapping and comparing immune landscapes rather than optimizing a single predictor, with more formal outcome modeling reserved for subsequent, targeted studies.

### Limitations

This study has several limitations that should temper interpretation. First, the mortality analyses rely on a single external transcriptomic cohort from the VANISH trial (Antcliffe *et al*. 2019), with a modest sample size and United Kingdom intensive-care setting; additional validation in larger, multi-center cohorts, different health systems, and non-septic critical illness will be needed to assess generalizability. Second, we focused on baseline, cross-sectional whole-blood expression and simple multivariable logistic regression, without modelling longitudinal trajectories, nonlinear relationships, or multi-omics integration, any of which might alter the relative performance of literature-derived versus data-driven signatures. Third, the semaglutide case study is anchored on serum proteomics from a single STEP trial (Maretty *et al*. 2025) and on proteins measurable by a SomaScan panel; membrane and intracellular signaling components of GLP-1 biology are incompletely captured, and we did not evaluate clinical infection endpoints, so our conclusions about infection-related pathways remain hypothesis-generating. Fourth, the LLM components inherit biases from both the underlying literature and the models themselves: PubMed-centered corpora under-represent negative and non-English studies, retrieval currently accesses only a small fraction of the embedding space, and behavior can vary with prompt design and model choice. Finally, our fine-tuned GPT-OSS judge was evaluated on semaglutide prompts without a gold-standard ranking or systematic comparison to expert adjudication, so it’s apparent calibration and concordance with the base model should be viewed as an initial behavioral characterization rather than definitive evidence of superiority.

## Conclusions

In summary, we extend a previously published LLM-guided, literature-based gene prioritization framework (Khan *et al*. 2025) to show that a compact sepsis priority set (PS3) can achieve independent mortality prediction in a randomized trial cohort, with performance comparable to larger data-driven immune signatures in the sickest patients. The same pipeline, when redirected toward GLP-1/obesity/infection biology, recapitulates the metabolic footprint of semaglutide and highlights immune–metabolic pathways relevant to infection, while revealing the assay-dependent nature of gene-versus protein-level concordance. Supervised fine-tuning of an open-weight LLM-as-judge on curated sepsis justifications preserves global ranking behavior yet induces interpretable shifts in immune and infection themes, suggesting a route to domain-aware evaluators that sit transparently on top of retrieval-driven screening. Together, these findings support the use of LLM-guided, literature-grounded gene sets as flexible, mechanistically interpretable modules for probing immune dysregulation across diseases and therapies, and they provide a template for integrating clinical validation, cross-domain case studies, and LLM-as-judge specialization in future biomarker development work.

## Reference

Antcliffe DB, Burnham KL, Al-Beidh F et al. Transcriptomic Signatures in Sepsis and a Differential Response to Steroids. From the VANISH Randomized Trial. Am J Respir Crit Care Med 2019;199:980–6.

Asgari E, Montaña-Brown N, Dubois M et al. A framework to assess clinical safety and hallucination rates of LLMs for medical text summarisation. npj Digit Med 2025;8:274.

Chen Q, Zhang M, Xia Y et al. Dynamic risk stratification and treatment optimization in sepsis: the role of NLPR. Front Pharmacol 2025;16:1572677.

Chenoweth JG, Brandsma J, Striegel DA et al. Sepsis endotypes identified by host gene expression across global cohorts. Commun Med 2024;4:120.

Davenport EE, Burnham KL, Radhakrishnan J et al. Genomic landscape of the individual host response and outcomes in sepsis: a prospective cohort study. The Lancet Respiratory Medicine 2016;4:259–71.

Diorio C, Shaw PA, Pequignot E et al. Diagnostic biomarkers to differentiate sepsis from cytokine release syndrome in critically ill children. Blood Adv 2020;4:5174–83.

Fang Z, Liu X, Peltz G. GSEApy: a comprehensive package for performing gene set enrichment analysis in Python. Bioinformatics 2023;39:btac757.

Farquhar S, Kossen J, Kuhn L et al. Detecting hallucinations in large language models using semantic entropy. Nature 2024;630:625–30.

Ganesan A, Moore AR, Zheng H et al. A conserved immune dysregulation signature is associated with infection severity, risk factors prior to infection, and treatment response. Immunity 2025;0, DOI: 10.1016/j.immuni.2025.05.020.

Hotchkiss RS, Monneret G, Payen D. Sepsis-induced immunosuppression: from cellular dysfunctions to immunotherapy. Nat Rev Immunol 2013;13:862–74.

Khan T, Toufiq M, Yurieva M et al. Automating candidate gene prioritization with large language models: from naive scoring to literature-grounded validation. Bioinformatics 2025;41:btaf541.

Kontsioti E, Maskell S, Pirmohamed M. Response to the comment on: Exploring the impact of design criteria for reference sets on performance evaluation of signal detection algorithms: The case of drug–drug interactions. Pharmacoepidemiology and Drug Safety 2024;33:e5731.

Kuleshov MV, Jones MR, Rouillard AD et al. Enrichr: a comprehensive gene set enrichment analysis web server 2016 update. Nucleic Acids Res 2016;44:W90–7.

Liberzon A, Birger C, Thorvaldsdóttir H et al. The Molecular Signatures Database (MSigDB) hallmark gene set collection. Cell Syst 2015;1:417–25.

Maretty L, Gill D, Simonsen L et al. Proteomic changes upon treatment with semaglutide in individuals with obesity. Nat Med 2025;31:267–77.

Moore AR, Zheng H, Ganesan A et al. A consensus immune dysregulation framework for sepsis and critical illnesses. Nat Med 2025:1–13.

Omar M, Sorin V, Collins JD et al. Multi-model assurance analysis showing large language models are highly vulnerable to adversarial hallucination attacks during clinical decision support. Commun Med 2025;5:330.

OpenAI, Agarwal S, Ahmad L et al. gpt-oss-120b & gpt-oss-20b Model Card. 2025, DOI: 10.48550/arXiv.2508.10925.

Pelaia TM, Shojaei M, McLean AS. The Role of Transcriptomics in Redefining Critical Illness. Crit Care 2023;27:89.

Qiu P, Wu C, Liu S et al. Quantifying the reasoning abilities of LLMs on clinical cases. Nat Commun 2025;16:9799.

Shool S, Adimi S, Saboori Amleshi R et al. A systematic review of large language model (LLM) evaluations in clinical medicine. BMC Med Inform Decis Mak 2025;25:117.

Singh A, D’Arcy M, Cohan A et al. SciRepEval: A Multi-Format Benchmark for Scientific Document Representations. arXiv, 2022.

Sparks R, Rachmaninoff N, Lau WW et al. A unified metric of human immune health. Nat Med 2024;30:2461–72.

Sweeney TE, Perumal TM, Henao R et al. A community approach to mortality prediction in sepsis via gene expression analysis. Nat Commun 2018;9:694.

Sweeney TE, Wong HR. Risk stratification and prognosis in sepsis: what have we learned from microarrays? Clin Chest Med 2016;37:209–18.

Taktaz F, Fontanella RA, Scisciola L et al. Bridging the gap between GLP1-receptor agonists and cardiovascular outcomes: evidence for the role of tirzepatide. Cardiovasc Diabetol 2024;23:242.

Toufiq M, Rinchai D, Bettacchioli E et al. Harnessing large language models (LLMs) for candidate gene prioritization and selection. J Transl Med 2023;16;21(1):728.

Transcriptomic Signatures in Sepsis and a Differential Response to Steroids. From the VANISH Randomized Trial | American Journal of Respiratory and Critical Care Medicine. https://www.atsjournals.org/doi/10.1164/rccm.201807-1419OC?url_ver=Z39.88-2003&rfr_id=ori:rid:crossref.org&rfr_dat=cr_pub%20%200pubmed (July 7, 2025, date last accessed)

Wong HR, Cvijanovich NZ, Anas N et al. Developing a clinically feasible personalized medicine approach to pediatric septic shock. Am J Respir Crit Care Med 2015;191:309–15.

Yugar LBT, Sedenho-Prado LG, da Silva Ferreira IMC et al. The efficacy and safety of GLP-1 receptor agonists in youth with type 2 diabetes: a meta-analysis. Diabetol Metab Syndr 2024;16:92.

Zhang C, Zhang X, Sun Z et al. MetaSepsisBase: a biomarker database for systems biological analysis and personalized diagnosis of heterogeneous human sepsis. Intensive Care Med 2023;49:1015–7.

Zheng H, Rao AM, Dermadi D et al. Multi-cohort analysis of host immune response identifies conserved protective and detrimental modules associated with severity across viruses. Immunity 2021;54:753-768.e5.

